# Automated assembly of a reference taxonomy for phylogenetic data synthesis

**DOI:** 10.1101/116418

**Authors:** Jonathan A. Rees, Karen Cranston

## Abstract

Taxonomy and nomenclature data are critical for any project that synthesizes biodiversity data, as most biodiversity data sets use taxonomic names to identify taxa. Open Tree of Life is one such project, synthesizing sets of published phylogenetic trees into comprehensive supertrees. No single published taxonomy met the taxonomic and nomenclatural needs of the project. Here we describe a system for reproducibly combining several source taxonomies into a synthetic taxonomy, and we discuss the challenges of taxonomic and nomenclatural synthesis for downstream biodiversity projects.

## Introduction

Any large biodiversity data project requires one or more taxonomies for discovery and data integration purposes, as in “find occurrence records for primates” or “find the taxon record associated with this sequence” (Page 2008). Examples of such projects are GBIF (Edwards 2004), which focuses on occurrence records, and NCBI (Federhen 2011), which focuses on genetic sequence records. Each of these projects has a dedicated taxonomy effort that is responsive to the project’s particular needs. We present the design and application of the Open Tree Taxonomy, which serves the Open Tree of Life project, an aggregation of phylogenetic trees with tools for operating on them.

In order to meet Open Tree’s project requirements, the taxonomy is an automated assembly of ten different source taxonomies. The assembly process is repeatable so that we can easily incorporate updates to source taxonomies. Repeatability also allows us to easily test potential improvements to the assembly method.

Information about taxa is typically expressed in databases and files in terms of taxon names or ‘name-strings’. To combine taxonomies it is therefore necessary to be able to determine name equivalence: whether or not an occurrence of a name-string in one data source refers to the same taxon as a given name-string occurrence in another. Solving this equivalence problem requires that we distinguishing occurrences that only coincidentally have the same name-string (homonym sense detection), and unify occurrences only when evidence justifies it. We have developed a set of heuristics that scalably address this equivalence problem.

### The Open Tree of Life project

The Open Tree of Life project consists of a set of tools for

1. synthesizing phylogenetic supertrees from a corpus of phylogenetic tree inputs (input trees)
2. matching groupings in supertrees with higher taxa (such as Mammalia)
3. supplementing supertrees with taxa obtained only from taxonomy.

The outcome is one or more summary trees combining phylogenetic and taxonomic knowledge. Fig. 1 illustrates an overview of the process of combining phylogeny and taxonomy, while the details are described in a separate publication (Redelings and Holder 2017). Although Open Tree is primarily a phylogenetics effort, it requires a reference taxonomy that can support each of these functions.

**Figure 1.**
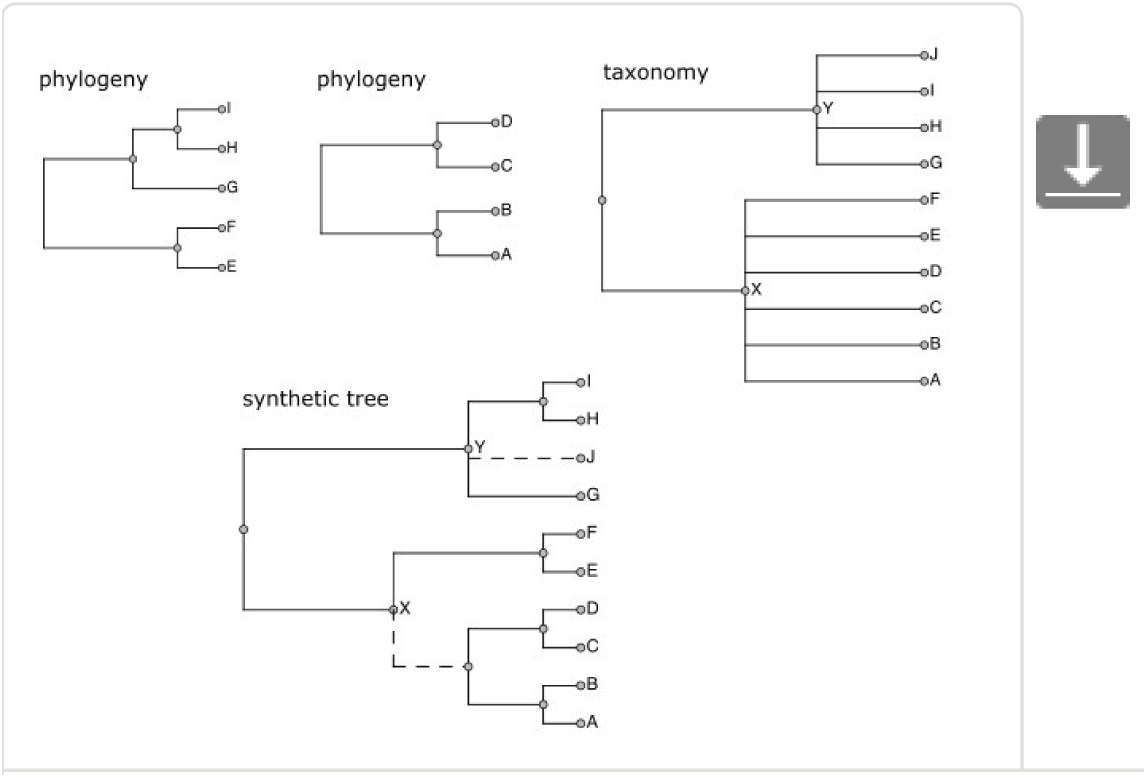
Role of taxonomy in assembly of the Open Tree of Life synthetic phylogeny. Dotted lines in the synthetic tree are those that come only from taxonomy, while solid lines have phylogenetic support. The taxonomy both links disjoint phylogenies and adds taxa not present in input trees.

For supertree synthesis (1), we use the taxonomy for converting OTUs (operational taxonomic units, or ‘tips’) on input trees to a canonical form. Supertree construction requires that a input tree OTU be matched with an OTU from another input tree whenever possible. This is a nontrivial task because a taxon can have very different OTU labels in different input trees due to synonymies, abbreviations, misspellings, notational differences, and so on. In addition, which taxon is named by a given label can vary across trees (homonymy). The approach we take is to map OTUs to the reference taxonomy, so that OTUs in different input trees are compared by comparing the taxa to which they map.

For higher taxon associations (2), we compare the groupings in the supertree to those in the taxonomy.

For supplementation (3), only a relatively small number of described taxa are represented in input trees (currently about 200,000 in the phylogenetic corpus out of two million or more known taxa), so the taxonomy provides those that are not. The large complement of taxonomy-only taxa can be ‘grafted’ onto a supertree in phylogenetically plausible locations based on how they relate taxonomically to taxa that are known from input trees.

### Reference taxonomy requirements

This overall program dictates what we should be looking for in a reference taxonomy. In addition to the technical requirements derived from the above, we have two additional requirements coming from a desire to situate Open Tree as ongoing infrastructure for the evolutionary biology community, rather than as a one-off study. Following are all five requirements:

1. **OTU coverage**: The reference taxonomy should have a taxon at the level of species or higher for every OTU that has the potential to occur in more than one study, over the intended scope of all cellular organisms.
2. **Phylogenetically informed classification**: Higher taxa should be provided with as much resolution and phylogenetic fidelity as is reasonable. Ranks and nomenclatural structure should not be required (since many well-established groups do not have proper Linnaean names or ranks) and groups at odds with phylogenetic understanding (such as Protozoa) should be avoided.
3. **Taxonomic coverage**: The taxonomy should cover as many as possible of the species that are described in the literature, so that we can supplement generated supertrees as described in step 3 above.
4. **Ongoing update**: New taxa of importance to phylogenetic studies are constantly being added to the literature. The taxonomy needs to be updated with new information on an ongoing basis.
5. **Open data**: The taxonomy must be available to anyone for unrestricted use. Users should not have to ask permission to copy and use the taxonomy, nor should they be bound by terms of use that interfere with further reuse.

An additional goal is that the process should be reproducible and transparent. Given the source taxonomies, we should be able to regenerate the taxonomy, and taxon records should provide information about the taxonomic sources from which it is derived.

No single available taxonomic source meets all requirements. The NCBI taxonomy has good coverage of OTUs, provides a rich source of phyogenetically informed higher taxa, and is open, but its taxonomic coverage is limited to taxa that have sequence data in GenBank (only about 360,000 NCBI species having standard binomial names at the time of this writing). Traditional all-life taxonomies such as Catalogue of Life (Roskov et al. 2016), IRMNG (Rees 2008), and GBIF meet the taxonomic coverage requirement, but miss many OTUs from our input trees, and their higher-level taxonomies are often not as phylogenetically informed or resolved as the NCBI taxonomy. At the very least, Open Tree needs to combine an NCBI-like sequence-aware taxonomy with a traditional broad taxonomy that is also open.

These requirements cannot be met in an absolute sense; each is a ‘best effort’ requirement subject to availability of project resources.

Note that the Open Tree Taxonomy is *not* supposed to be a reference for nomenclature; it links to other sources for nomenclatural and other information. Nor is it a place to deposit curated taxonomic information. The taxonomy has not been vetted in detail, as doing so would be beyond the capacity and focus of the Open Tree project. It is known to contain many taxon duplications and technical artifacts. Tolerating these shortcomings is a necessary tradeoff in attempting to meet the above requirements.

## Method

The conventional approach to meeting the requirements stated in the introduction would be to create a database, copy the first taxonomy into it, then somehow merge the second taxonomy into that, repeating for further sources if necessary. However, it is not clear how to meet the ongoing update requirement under this approach. As the source taxonomies change, we would like for the combined taxonomy to contain only information derived from the latest versions of the sources, without residual information from previous versions. Many changes to the sources are corrections, and we do not want to retain information from previous versions that is known to be incorrect.

Rather than maintain a database of taxonomic information, we instead developed a process for assembling a taxonomy from two or more taxonomic sources. With a repeatable process, we can generate a new combined taxonomy version from new source taxonomy versions *de novo,* and do so frequently. There are additional benefits as well, such as the ability to add new sources relatively easily, and to use the tool for other purposes.

In the following, any definite claims or measurements refer to the Open Tree reference taxonomy version 3.0.

### Terminology

- source taxonomy = imported taxonomic source (NCBI taxonomy, etc.)
- workspace = data structure for creation of the reference taxonomy
- name-string = taxonomic name considered textually, without association with any particular taxon or nomenclatural code
- node = a taxon record, either from a source taxonomy or the workspace. Records primary name-string, provenance, parent node, optional rank, optional annotations
- parent (node) = the nearest enclosing node within a given node’s taxonomy
- tip = a node that is not the parent of any node
- homonym = where a single name-string belongs to multiple nodes within the same taxonomy. This is close to the nontechnical meaning of ‘homonym’ and is not to be confused with ‘homonym’ in the nomenclatural sense, which only applies within a single nomenclatural code. Nomenclatural homonyms and hemihomonyms (Shipunov 2011) are both homonyms in this sense.
- primary = the non-synonym name-string of a node, as opposed to one of the synonyms.
- image (of a node n’) = the workspace node corresponding to n’
- *incertae sedis:* taxon A is *incertae sedis* in taxon B if A is a child of B but is not known to be disjoint from B’S non*-incertae-sedis* children. That is, if we had more information, it might turn out that A is a member of one of the other children of B.

### Method overview

This section is an overview of the taxonomy assembly method. Several generalities stated here are simplifications; the actual method (described later) is significantly more involved.

We start with a sequence of source taxonomies S1, S2, …, Sn, ordered by priority. Priority means that if S is judged more accurate or otherwise “better” than S’, then S occurs earlier in the sequence than S’ and its information supersedes that from later sources. Priority judgements are made by curators (either project personnel or participants in Open Tree workshops and online forums) based on their taxonomic expertise.

We define an operator for combining taxonomies pairwise, written schematically as U = S + S’, and apply it from left to right:

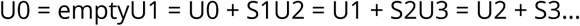

The combination S + S’ is formed in two steps:

1. A mapping or *alignment* step that identifies all nodes in S’ that can be equated with nodes in S. There will often be nodes in S’ that cannot be aligned to S.
2. A *merge* step that creates the combination U = S + S’, by adding to S the unaligned taxa from S’. The attachment position of unaligned nodes from step 1 is determined from nearby aligned nodes, either as a *graft* or an *insertion.*

Examples of these two cases are given in Figure 2.

AS a simple example, consider a genus represented in both taxonomies, but containing different species in the two:

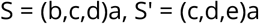

S and S’ each have four nodes. Suppose c, d, and a in S’ are aligned to c, d, and a in S. The only unaligned node is e, which is a sibling of c and d and therefore grafted as a child of a. After the merge step, we have:

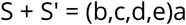

One might call this merge heuristic ‘my sibling’s sibling is my sibling’ or ‘transitivity of siblinghood’.

This is a very common pattern. Fig. 2 illustrates a real life-example when combining the genus *Bufo* across NCBI and GBIF. There are about 900,000 similar simple grafting events in the assembly of OTT.

**Figure 2.**
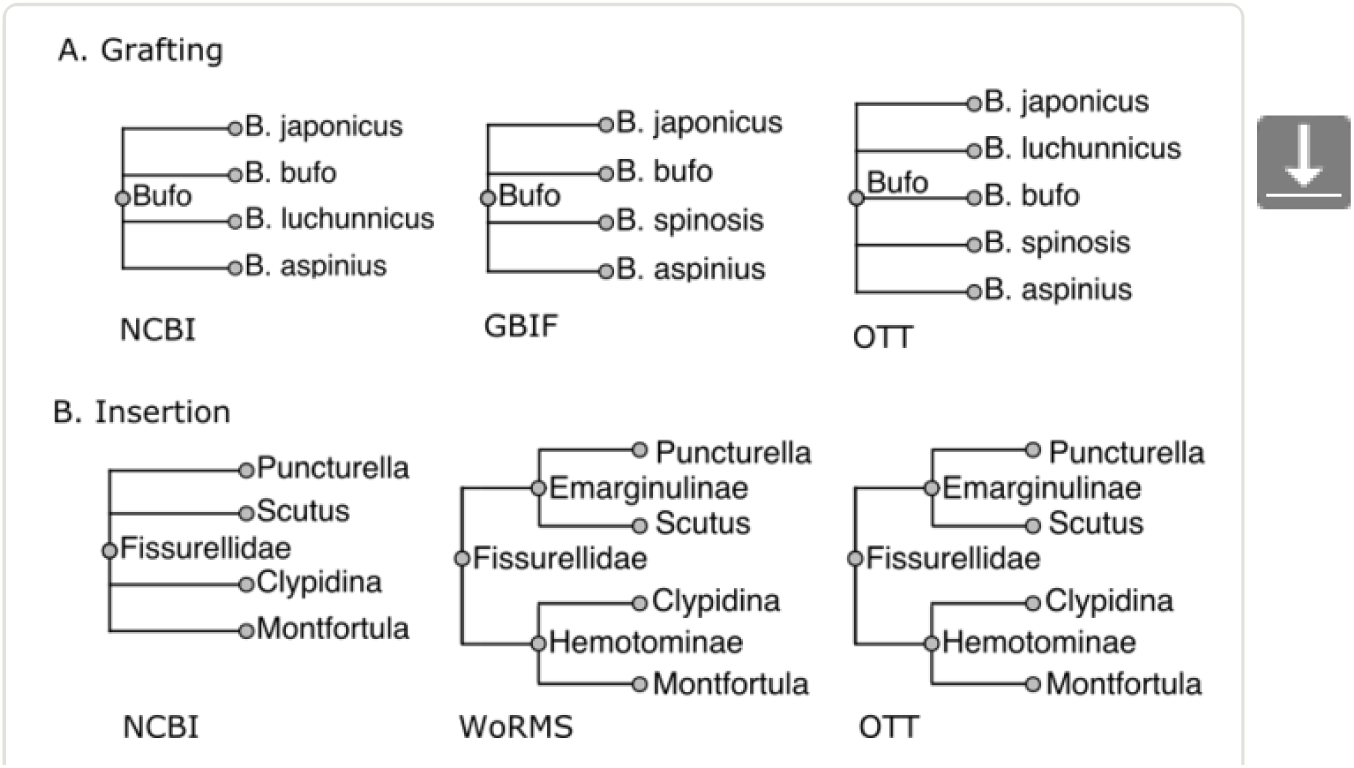
Examples of grafting and insertion when combining taxonomies. In both cases, the NCBI taxonomy has higher priority than GBIF. In A (grafting), we assemble the genus *Bufo* across NCBI and GBIF. There is no *B. spinosis* in GBIF and no *B. luchunnicus* in NCBI. Therefore, the *Bufo* in the combined taxonomy has as its children copies of species records from both sources. In B (insertion), WoRMS provides greater resolution of *Fissurellidae* than NCBI taxonomy: it divides the family into subfamilies *Hemotominae* and *Emarginulinae,* nodes that do not exist in NCBI. The subfamilies are ‘inserted’ in a way that adds information without disrupting existing relationships from NCBI.

The other merge method is an *insertion,* where the unaligned node has descendants that are in S. This always occurs when S’ has greater resolution than S. For example, see Fig. 2, where WoRMS provides greater resolution than NCBI.

The vast majority of alignment and merge situations are simple, similar to the examples shown in Fig. 2. However, even a small fraction of special cases can add up to thousands when the total number of alignments and merges measures in the millions, so we have worked to develop heuristics that handle the most common special cases. Ambiguities caused by synonyms and homonyms create most of the difficulties, with inconsistent or unclear higher taxon membership creating the rest. The development of the assembly process described here has been a driven by trial and error - finding cases that fail and then adding / modifying heuristics to address the underlying cause, or adding an *ad hoc* adjustment for cases that are rare or overly complex.

### Taxonomic sources

We build the taxonomy from ten sources. Some of these sources are from taxonomy projects, while others were manually assembled based on recent publications. OTT assembly is dependent on the input order of the sources - higher ranked inputs take priority over lower ranked inputs. Table 1 lists the sources used to construct OTT. The full provenance details, and a copy of the normalized source, are available in supplementary data.

**Table 1.**
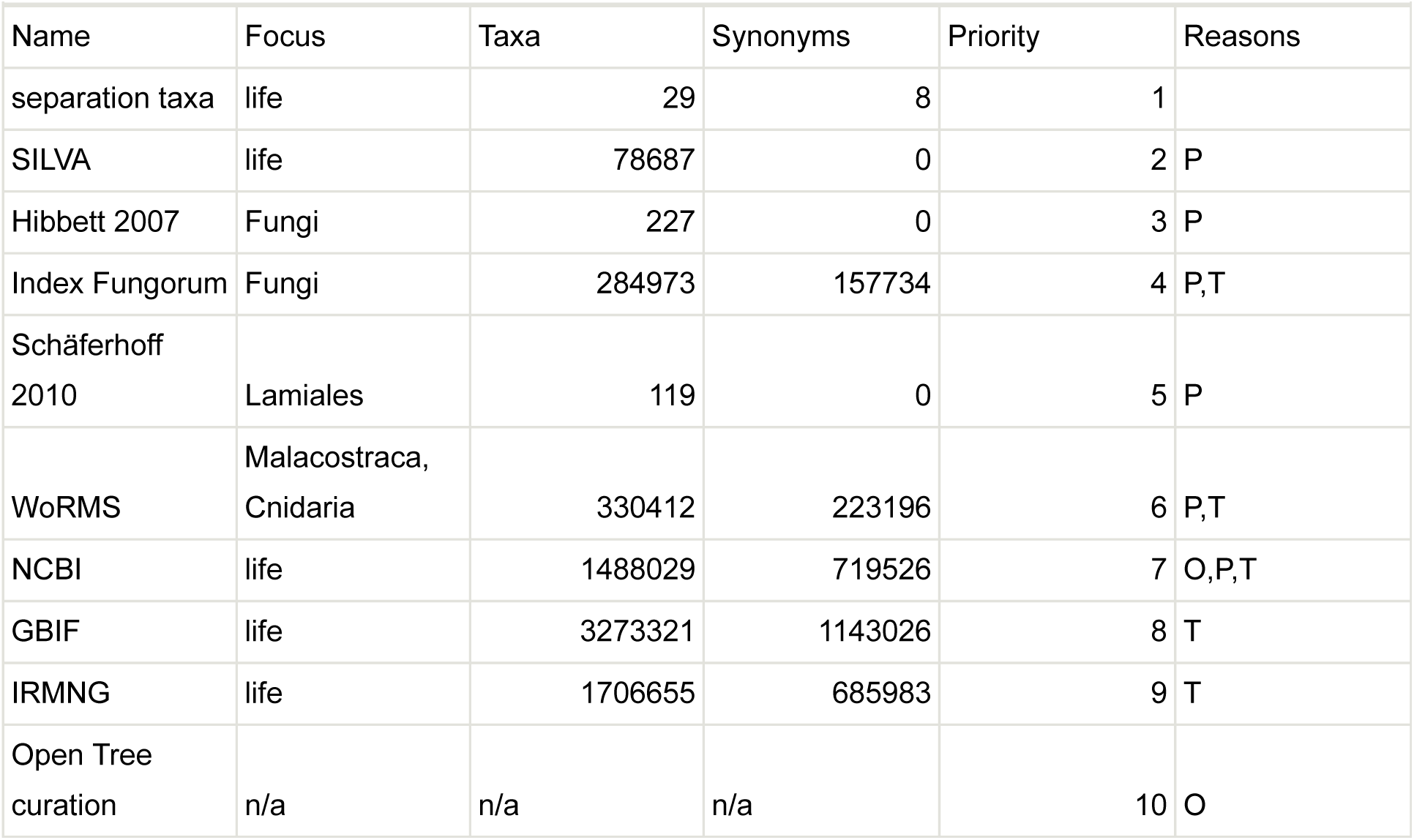
Download as CSV. List of taxonomic sources: The ten sources used in v3.0 of Open Tree Taxonomy. Six are online taxonomic resources (SILVA (Quast et al. 2013), Index Fungorum (Index Fungorum Partnership 2014b), WoRMS (WORMS Editorial Board 2015), NCBI, GBIF, IRMNG), two are from publications (Hibbett et al. 2007,Schäferhoff et al. 2010), one is a small curated taxonomy of major groups to aid in assembly (“separation taxa”), and one consisting of taxonomic additions from phylogenies input into the Open Tree system (“Open Tree curation”). See text for explanation of ‘Open Tree curation’ and ‘separation taxa’. Detailed provenance information for each source can be found in the accompanying data package. Key to ‘reasons’ column: O = added in order to improve OTU coverage; P = added in order to improve phylogenetic classification; T = added in order to improve taxonomic coverage.

| Name | Focus | Taxa | Synonyms | Priority | Reasons |
| --- | --- | --- | --- | --- | --- |
| separation taxa | life | 29 | 8 | 1 |  |
| SILVA | life | 78687 | 0 | 2 | P |
| Hibbett 2007 | Fungi | 227 | 0 | 3 | P |
| Index Fungorum | Fungi | 284973 | 157734 | 4 | P,T |
| Schäferhoff<br>2010 | Lamiales | 119 | 0 | 5 | P |
| WoRMS | Malacostraca,<br>Cnidaria | 330412 | 223196 | 6 | P,T |
| NCBI | life | 1488029 | 719526 | 7 | O,P,T |
| GBIF | life | 3273321 | 1143026 | 8 | T |
| IRMNG | life | 1706655 | 685983 | 9 | T |
| Open Tree<br>curation | n/a | n/a | n/a | 10 | O |

#### Open Tree curation

It is not uncommon to have taxa as OTUs in phylogenetic studies that do not occur in OTT. This can be due to a delay in curation by the source taxonomy, a delay in importing a fresh source version into OTT, a morphological study containing otherwise unknown species, or other causes. To handle this situation, we developed a user interface that allows curators to create new taxon records along with relevant documentation (publications, databases, and so on). New taxon records are saved into a public GitHub repository, and these records are then linked from the OTT taxonomy files and user interfaces so that provenance is always available.

#### Separation taxa

This is a small curated tree containing 31 major groups such as Fungi, Metazoa, and Lepidoptera. Its purpose is to assist in separating homonyms. If a node is found in one of these separation groups, then it will not match a node in a disjoint separation group, absent other evidence (details below).

#### ARB-SILVA taxonomy processing

The terminal taxa in the SILVA taxonomy are algorithmically generated clusters of RNA sequences derived from GenBank records. Rather than incorporate these idiosyncratic, fine-grained groupings into OTT, we use sequence record metadata to group the clusters into larger groups corresponding to NCBI taxa, and include those larger groups in OTT.

We excluded SILVA’s plant, animal, and fungal branches from OTT because these groups are well covered by other sources and poorly represented in SILVA. For example, SILVA has only 299 taxa in Metazoa, compared with over 500,000 taxa under Metazoa in NCBI Taxonomy.

#### Extinct / extant annotations

Curators requested information about whether taxa were extinct vs. extant. With the exception of limited data from WoRMS and Index Fungorum, this information was not explicitly present in our other sources, so we imported IRMNG, which logs the extinct / extant status of taxa.

As a secondary heuristic, records from GBIF that originate from PaleoDB, and do not come from any other taxonomic source, are annotated extinct. This is not completely reliable, as some PaleoDB taxa are extant.

#### Suppressed records

We suppress the following source taxonomy records:

- animals, plants, fungi in SILVA
- GBIF backbone records that originate from IRMNG (IRMNG is imported separately)
- GBIF backbone records that originate from IPNI
- GBIF backbone records whose taxonomic status is ‘doubtful’
- GBIF backbone records for infraspecific taxa (subspecies, variety, form)
- IRMNG records whose nomenclatural status is ‘nudum’, ‘invalid’, or any of about 25 similar designations
- NCBI Taxonomy records that cannot correspond to taxa: those with names containing ‘uncultured’, ‘unidentified’, ‘insertion sequences’, or any of about 15 similar designations

The IPNI and IRMNG derived GBIF records are suppressed because they include many invalid names. We pick up most of the valid names from other sources, such as the direct IRMNG import, so this is not a great loss. Although GBIF’s original taxonomic sources indicate which names are known to be invalid, this information is not provided by the GBIF backbone. Note that the GBIF backbone might import the same name from more than one source, but its provenance information only lists one of the sources. We suppress the record if that one source is IPNI or IRMNG.

### Import and Normalization

Each source taxonomy has its own import procedure, usually a file download from the provider’s web site followed by application of a script that converts the source to a common format for import. Given the converted source files, the taxonomy can be read by the OTT assembly procedure.

After each source taxonomy is loaded, the following normalizations are performed:

1. Diacritics removal - accents, umlauts, and other diacritic marks are removed in order to improve name matching, as well as to follow the nomenclatural codes, which prohibit them. The original name-string is kept as a synonym.
2. Child taxa of “containers” in the source taxonomy are made to be children of the container’s parent. “Containers” are groupings in the source that don’t represent taxa, for example nodes named “incertae sedis” or “environmental samples”. The members of a container aren’t more closely related to one another than they are to the container’s siblings; the container is only present as a way to say something about the members. The fact that a node had originally been in a container is recorded as a flag on the child node.
3. When a subgenus X has the same name-string as its containing genus, its name-string is changed to “X subgenus X”. This follows a convention used by NCBI Taxonomy and helps distinguish the two taxa later in assembly.

The normalized versions of the taxonomies then become the input to subsequent processing phases.

### Aligning nodes across taxonomies

This section and the next give details of the taxonomy combination method introduced above.

OTT is assembled in a temporary work area or *workspace* by alternately aligning a source to the workspace and merging that source into the workspace. It is important that source taxonomy nodes be matched with workspace nodes when and only when this is appropriate. A mistaken identity between a source node and a workspace node can be disastrous, leading not just to an incorrect classification but to downstream curation errors in OTU matching (e.g. putting a snail in flatworms). A mistaken non-identity (separation) can also be a problem, since taxon duplication (i.e. multiple nodes for the same taxon) leads to loss of unification opportunities in phylogeny synthesis.

As described above, source taxonomies are processed (aligned and merged) in priority order. For each source taxonomy, *ad hoc* adjustments are applied before automatic alignments. For automatic alignment, alignments closest to the tips of the source taxonomy are found in a first pass, and all others in a second pass. The two-pass structure permits first-pass alignments to be used during the second pass (see Overlap, below).

#### Ad hoc adjustments

A set of *ad hoc* ‘adjustments’ address known issues that are beyond the capabilities of the automated process to address. These often reflect either errors or missing information in source taxonomies, discovered through the failure of automated alignment, and confirmed manually via the literature. Although each individual adjustment is *ad hoc,* i.e. not the result of automation, the adjustments are recorded in a file that can be run as a script. Following are some examples of adjustments.

1. capitalization and spelling repairs (e.g. change ‘sordariomyceta’ to ‘Sordariomyceta’)
2. addition of synonyms to facilitate later matching (e.g. ‘Florideophyceae’ as synonym for ‘Florideophycidae’)
3. name changes (e.g. ‘Choanomonada’ to ‘Choanoflagellida’)
4. deletions (e.g. removing synonym ‘Eucarya’ for ‘Eukaryota’ to avoid confusing eukaryotes with genus *Eucarya* in Magnoliopsida; or removing unaccepted genus *Tipuloidea* in Hemiptera to avoid confusion with the superfamily in Diptera)
5. merges to repair redundancies in the source (e.g. Pinidae, Coniferophyta, Coniferopsida)
6. rename taxa to avoid confusing homonyms (e.g. there are two Cyanobacterias in SILVA, one a parent of the other; the parent is renamed to its NCBI name ‘Cyanobacteria/Melainabacteria group’)
7. alignments when names differs (Diatomea is Bacillariophyta)
8. alignments to override automated alignment rules (Eccrinales not in Fungi, Myzostomatida not in Annelida)

In the process of assembling the reference taxonomy, about 300 *ad hoc* adjustments are made to the source taxonomies before they are aligned to the workspace.

#### Candidate identification

Given a source node, the alignment procedure begins by finding the nodes in the workspace that it could *possibly* align with. These workspace nodes are called *candidates.* The candidates are simply the nodes that have a name-string (either primary or synonym) that matches any name-string (primary or synonym) of the source node.

Example: GBIF *Nakazawaea pomicola* has NCBI *Candida pomiphila* as a candidate by way of an NCBI record that lists *Nakazawaea pomicola* as a synonym of *Candida pomiphila.*

It follows that if the workspace has multiple nodes with the same name-string (homonyms), all of these nodes will become candidates for every source node that also has that name-string.

#### Candidate selection

The purpose of the alignment phase is to choose a single correct candidate for each source node, or to reject all candidates if none is correct. For over 97% of source nodes, there are no candidates or only one candidate, and selection is fairly simple, but the remaining nodes require special treatment.

Example: There are two nodes named *Aporia lemoulti* in the GBIF backbone taxonomy; one is a plant and the other is an insect. One of these two is an erroneous duplication, but the automated system has to be able to cope with this situation because we don’t have the resources to correct all source taxonomy errors. When IRMNG is aligned, it is necessary to choose the right candidate for the node with name *Aporia lemoulti.* Consequences of incorrect placement might include putting siblings of IRMNG *Aporia lemoulti* in the wrong kingdom as well.

Example: *Fritillaria messanensis* in WoRMS must not map to *Fritillaria messanensis* in NCBI Taxonomy because the taxon in WoRMS is an animal (tunicate) while the taxon in NCBI is a flowering plant. This is a case where there is a unique candidate, but it is wrong.

Similarly, *Aporia sordida* is a plant in GBIF, but an insect in IRMNG.

#### Alignment heuristics

Once we have a list of candidates, we apply a set of heuristics in an attempt to find a single candidate, and thereby align a source node n’ with a workspace node n. The heuristics are as follows, presented in the order that we apply them in the alignment process:

1. **Separation**: If n and n’ are contained in “obviously different” major groups such as animals and plants, do not align n’ to n. Two major groups (or “separation taxa”) are “obviously different” if they are disjoint as determined by the separation taxonomy. *Examples:* (1) the *Aporia* cases above; (2) NCBI says n = *Pteridium* is a land plant, WoRMS says n’ = *Pteridium* is a rhodophyte, and the separation taxonomy says land plants and rhodophytes are disjoint, so n and n’ are different taxa.
2. **Disparate ranks**: Prohibit alignment where n and n’ have “obviously incompatible” (disparate) ranks. A rank is “obviously incompatible” with another if one is genus or a rank inferior to genus (species, etc.) and the other is family or a rank superior to family (order, etc.). *Examples:* (1) IRMNG *Pulicomorpha,* a genus, matches NCBI *Pulicomorpha,* a genus, not GBIF Pulicomorpha, a suborder. Note that both candidates are insects. (2) For genus *Ascophora* in GBIF (which is in Platyhelminthes), candidate *Ascophora* from WoRMS, a genus, is preferred to candidate Ascophora from NCBI, an infraorder.
3. **Lineage**: Prefer to align species or genus n’ to n if they have common lineage. For example, if n’ is a species, prefer candidates n where the name-string of the family-rank ancestor node of n’ is the same as the name-string of the family-rank ancestor node of n. *Example:* Source node *Plasmodiophora diplantherae* from Index Fungorum, in Protozoa, has one workspace candidate derived from NCBI and another from WoRMS. Because the source node and the NCBI candidate both claim to be in a taxon with name ‘Phytomyxea’, while the WoRMS candidate has no near lineage in common, the NCBI candidate is chosen. The details are complicated because (a) every pair of nodes have at least *some* of their lineage in common, and (b) genus names do not provide any information when comparing species nodes with the same name-string, so for species we can’t just look at the parent taxon. The exact rule used is the following: Define the ‘quasiparent name’ of n, q(n), to be the name-string of the nearest ancestor of n whose name-string is not a prefix of n’s name-string. (For example, the quasiparent of a species would typically be a family.) If q(n) is the name-string of an ancestor of n’, or q(n’) is the name-string of an ancestor of n, then prefer n to candidates that lack these properties.
4. **Overlap**: Prefer to align n’ to n if they are higher level groupings that overlap. Stated a bit more carefully: Prefern’ if n’ has a descendant aligned to a descendant of n. *Example:* Source node *Peranema* from GBIF has two candidates from NCBI. One candidate shares descendant *Peranema cryptocercum* with the source taxon, while the other shares no descendants with the source taxon. The source is therefore aligned to the one with the shared descendant.
5. **Proximity**: Require a candidate n to have the property that the smallest separation taxon containing the source node n’ is also the smallest separation taxon containing n. *Example:* for source node Heterocheilidae in IRMNG (a nematode family) whose smallest separation ancestor is Metazoa, choose the workspace (NCBI) candidate with smallest separation ancestor Metazoa (also a nematode family), and not the one with smallest separation ancestor Diptera (a fly family).
6. **Same name-string**: Prefer candidates whose primary name-string is the same as the primary name-string of n’. *Example:* For source node n’ = GBIF *Zabelia tyaihyoni,* candidate *Zabelia tyaihyoni* from NCBI is preferred to candidate *Zabelia mosanensis,* also from NCBI. NCBI *Z. mosanensis* is a candidate for n’ because GBIF declares that *Z. mosanensis* is a synonym for GBIF *Z. tyaihyoni.*

#### Control flow for applying heuristics

Each heuristic, when presented with a source node and a candidate (workspace node), answers ‘yes’, ‘no’, or ‘no information’. ‘Yes’ means that according to the rule, the two nodes refer to the same taxon, ‘no’ means they refer to different taxa, and ‘no information’ means that this rule provides no information as to whether the nodes refer to the same taxon.

The answers are assigned numeric scores of 1 for yes, 0 for no information, and -1 for no. A candidate that a heuristic gives a no is eliminated, one that is unique in getting a yes is selected, and if there are no yeses or no unique yes, more heuristics are consulted.

The heuristics are applied in the order in which they are listed above. The outcome is sensitive to the ordering. The ordering is forced to some extent by internal logic, but overall the ordering was determined by trial and error.

If there is a single candidate that is not rejected by any heuristic, it is aligned to that candidate.

More specifically, the method for applying the heuristics is as follows:

1. Start with a source node N and its set C of workspace node candidates C1 … Cn.
2. For each heuristic H as listed above:

1. For each candidate Ci currently in C, use H to obtain the score H(N, Ci)
2. Let Z = the highest score from among the scores H(N, Ci)
3. If Z < 0, we are done - no candidate is suitable
4. Let C’ = those members of C that have score Z
5. If Z > 0 and C’ contains only one candidate, we are done (match is that candidate)
6. Otherwise, replace C with C’ and proceed to the next heuristic
3. If C is singleton after all heuristics are exhausted, its member is taken to be the correct match.
4. Otherwise, the source node does not match unambiguously, and alignment fails.

#### Failure to choose

If the alignment process ends with multiple candidates, there is an unresolvable ambiguity. If the ambiguous source node has no children, it is dropped, which is OK because it probably corresponds to one of the existing candidates and therefore would make no new contribution. If the ambiguous source node has children, it is treated as unaligned and therefore new, possibly turning an N-way homonym into an N+1-way homonym. This could easily be wrong because it is so unlikely that the source node really represents a distinct taxon. Usually, the subsequent merge phase determines that the grouping is not needed because it inconsistent or can be ‘absorbed’, and it is dropped. If it is not dropped, then there is a troublesome situation that calls for manual review.

As an example of an unaligned tip, consider GBIF *Katoella pulchra.* The candidates are NCBI *Davallodes pulchra* and *Davallodes yunnanensis.* (There is no *Katoella pulchra* in the workspace at the time of alignment. The two candidates come from synonymies with *Katoella pulchra* declared by GBIF.) Neither candidate is preferable to the other, so *Katoella pulchra* is left unaligned and is omitted from the assembly.

## Merging unaligned source nodes

After the alignment phase, we are left with the set of source nodes that could not be aligned to the workspace. The next step is to determine if and how these (potentially new) nodes can be merged into the workspace.

The combined taxonomy (U, above) is constructed by adding copies of unaligned nodes from the source taxonomy S’ one at a time to the workspace, which initially contains a copy of S. Nodes of S’ therefore correspond to workspace nodes in either of two ways: by mapping to a copy of an S-node (via the S’-S alignment), or by mapping to a copy of an S’-node (when there is no S’-S alignment for the S’-node).

As described above, each copied S’-node is part of either a graft or an insertion. A graft or insertion rooted at r’ is attached to the workspace as a child of the nearest common ancestor node of r’’s siblings’ images. A graft is flagged *incertae sedis* if that NCA is a node other than the parent of the sibling images. By construction, insertions, never have this property, so an insertion is never flagged *incertae sedis*.

The following schematic examples illustrate each of the cases that come up while merging taxonomies. Taxonomy fragments are written in Newick notation (Olsen 1990). Fig. 3 illustrates each of these six cases.

**Figure 3.**
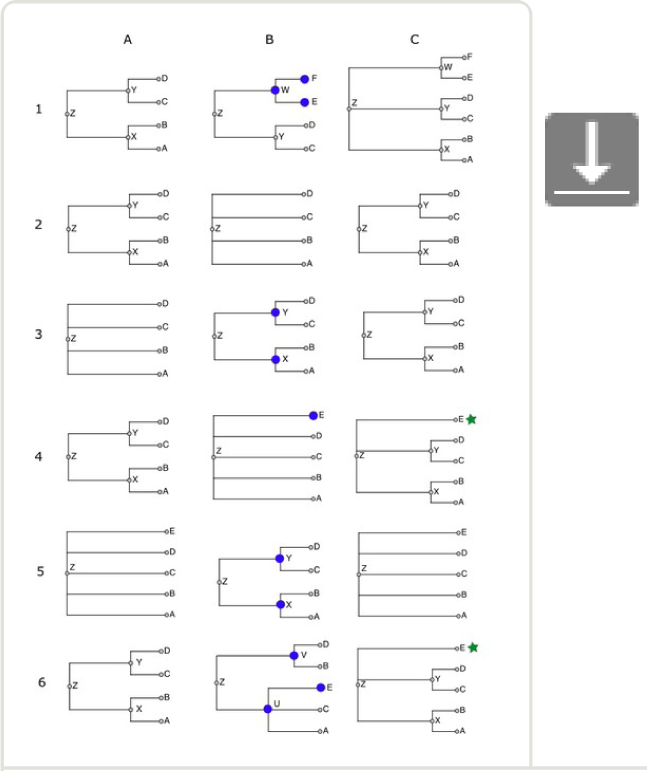
Merging taxonomies: Examples of outcomes when merging nodes from a source taxonomy into the workspace taxonomy. Each row (1-6) corresponds to one of the six examples described in the text. Column A is the current workspace taxonomy, column B is the source taxonomy being merged, and column C is the resulting workspace taxonomy. Nodes in B marked with a large blue circle are those that cannot be aligned to the workspace. Nodes in C marked with a green star are those flagged as *incertae sedis* in the final taxonomy.

### Case 1: ((a,b)x,(c,d)y)z + ((c,d)y,(e,f)w)z = ((a,b)x,(c,d)y,(e,f)w)z

This is a simple graft. The taxon w does not occur in the workspace, so it and its children are copied. The workspace copy of w is attached as a sibling of its siblings’ images: its sibling is y in S’, which is aligned to y in the workspace, so the copy becomes a child of y’s parent, or z.

### Case 2: ((a,b)x,(c,d)y)z + (a,b,c,d)z = ((a,b)x,(c,d)y)z

No nodes are copied from S’ to the workspace because every node in S’ is aligned to some node in S - there are no nodes that *could* be copied.

### Case 3:(a,b,c,d)z + ((a,b)x,(c,d)y)z = ((a,b)x,(c,d)y)z

Supposing x and y are unaligned, then x and y from S’ insert into the classification of z. The workspace gets copies of these two S’-nodes.

Example: superfamily Chitonoidea, which is in WoRMS (S’) but not in NCBI Taxonomy (S), inserts into NCBI Taxonomy. Its parent is suborder Chitonina, which is in NCBI (i.e. aligned to the workspace), and its children are six families that are all in NCBI (aligned).

### Case 4: ((a,b)x,(c,d)y)z + (a,b,c,d,e)z = ((a,b)x,(c,d)y,?e)z

In this situation, we don’t know where to put the unaligned taxon e from S’: in x, in y, or in z (sibling to x and y). The solution used here is to add e to z and mark it as *incertae sedis* (indicated above by the question mark).

For example, family Melyridae from GBIF has five genera, of which two (*Trichoceble, Danacaea*) are not found in the workspace, and the other three do not all have the same parent after alignment - they are in three different subfamilies. *Trichoceble* and *Danacaea* are made to be *incertae sedis* children of Melyridae, because there is no telling which NCBI subfamily they are supposed to go in.

### Case 5: (a,b,c,d,e)z + ((a,b)x,(c,d)y)z = (a,b,c,d,e)z

We don’t want to lose the fact from the higher priority taxonomy S that e is a proper child of z (i.e. not *incertae sedis*), so we discard nodes x and y, ignoring what would otherwise have been an insertion.

So that we have a term for this situation, say that x is *absorbed* into z.

### Case 6: ((a,b)x,(c,d)y)z + ((a,c)p,(b,d,e)q)z = ((a,b)x,(c,d)y,?e)z

If the source has a hierarchy that is incompatible with the one in the workspace, the conflicting source nodes are ignored, and any unaligned nodes (e) become *incertae sedis* nodes under an ancestor containing the incompatible node’s children.

For example, when WoRMS is merged, the workspace has, from NCBI,

((Archaeognatha)Monocondylia,(Pterygota,Zygentoma)Dicondylia)Insecta and the classification given by WoRMS is

((Archaeognatha,Thysanura=Zygentoma)Apteryogota,Pterygota)Insecta

That is, NCBI groups Thysanura (Zygentoma) with Pterygota, while WoRMS groups it with Archaeognatha. The WoRMS hierarchy is ignored in favor of the higher priority NCBI hierarchy. If Insecta in WoRMS had had an unaligned third child, it would have ended up *incertae sedis* in Insecta.

The test for compatibility is very simple: a source node is incompatible with the workspace if the nodes that its aligned children align with do not all have the same parent.

## Final patches

After all source taxonomies are aligned and merged, we apply general *ad hoc* additions and patches to the workspace, in a manner similar to that employed with the source taxonomies. Patches are represented in three formats. An early patch system used hand-written tabular files, additions via the user interface use a machine-processed JSON format, and most other patches are written as simple Python statements. There are 106 additions in JSON form, 97 additions and patches in tabular form, and approximately 121 in Python form.

## Assigning identifiers

The final step is to assign unique, stable identifiers to nodes so that external links to OTT nodes will continue to function correctly after the previous OTT version is replaced by the new one.

Identifier assignment is done by aligning the previous version of OTT to the new version. As with the other alignments, there are scripted *ad hoc* adjustments to correct for some errors that would otherwise be made by automated assignment. For this alignment, the set of heuristics is extended by adding rules that prefer candidates that have the same source taxonomy node id as the previous version node being aligned. After transferring identifiers of aligned nodes, any remaining workspace nodes are given newly ‘minted’ identifiers.

The alignment is computed only for the purpose of assigning identifiers; the previous OTT version is not merged into the workspace. An identifier can only persist from one OTT version to the next if it continues to occur in some source taxonomy.

## Results

The assembly method described above yields the reference taxonomy that is used by the Open Tree of Life project. The taxonomy itself, the details of how the assembly method unrolls to generate the taxonomy, and the degree to which the taxonomy meets the goals set out for it are all of interest in assessing how, and how well, the method works. We will address each of these three aspects of the method in turn.

## Summary of Open Tree Taxonomy

The methods and results presented here are for version 3.0 of the Open Tree Taxonomy (which follows five previous releases using the automated assembly method). The taxonomy contains 3,594,550 total taxa; 3,272,177 tips; and 277,365 internal nodes. 2,335,412 of the nodes have a Linnean binomial of the form *Genus epithet.* There are 1,842,403 synonym records and 9,089 homonyms (name-strings for which there are multiple nodes). A longer list of metrics is in Table 2.

**Table 2.** Download as CSV. Summary of Open Tree Taxonomy 3.

| Number nodes | Property |
| --- | --- |
| 3594550 | Total taxon records (nodes) |
| 1842403 | Synonym records |
| 277365 | Internal (non-tip) nodes |
| 3272177 | Tips. |
| 3116485 | Rank of 'species' |
| 70886 | Below the rank of species (e.g. subspecies, variety) |
| 67070 | Above the rank of species that subtend no node of rank species |
| 2335412 | Name-string has the form of a Linnaean binomial Genus epithet |
| 9089 | Homonym name-strings |
| 2867 | Homonym name-strings where the nodes have species rank |
| 6110 | Homonym name-strings where the nodes have genus rank |
| 38 | Maximum nesting depth of any node in the taxonomy |
| 53287 | Maximum number of children for any node in the taxonomy |
| 12.96 | Branching factor (average number of children per internal node) |
| 45008 | Source taxa that were absorbed into a larger taxon |
| 317624 | Marked *incertae sedis* or equivalent |
| 252600 | Annotated as being for an extinct taxon |

## Results of assembly procedure

As OTT is assembled, the alignment procedure processes every source node, either choosing an alignment target for it in the workspace based on the results of the heuristics, or leaving it unaligned. Fig. 4 illustrates the action of the alignment phase. The presence of a single candidate node does not automatically align the two nodes - we still apply the heuristics to ensure a match (and occasionally reject the single candidate).

**Figure 4.**
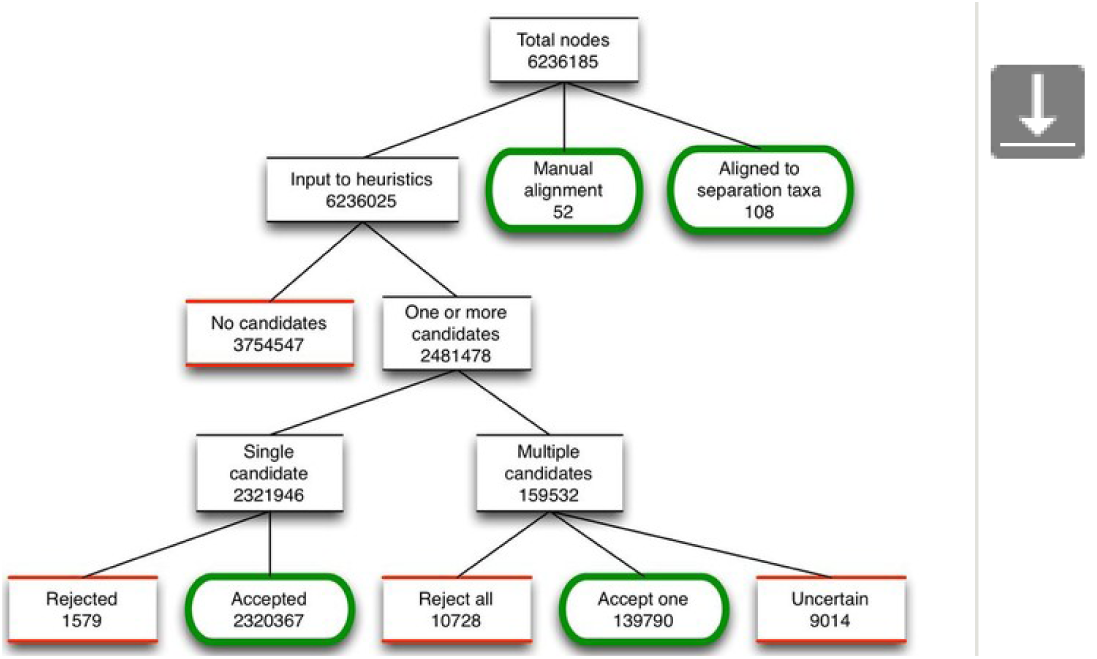
Fate of nodes as they move through the alignment procedure. Green, rounded boxes are endpoints that result in aligned nodes, while red, sqaure boxes are endpoints that result in unaligned nodes.

We counted the frequency of success for each heuristic, i.e. the number of times that a particular heuristic was the one that accepted the winning candidate from among two or more candidates. Table 3 shows these results. Separation (do not align taxa in disjoint separation taxa; used first), Lineage (align taxa with shared lineage; used midway through) and Same-name-string (prefer candidates who primary name-string matches; used last) were by far the most frequent.

**Table 3.**
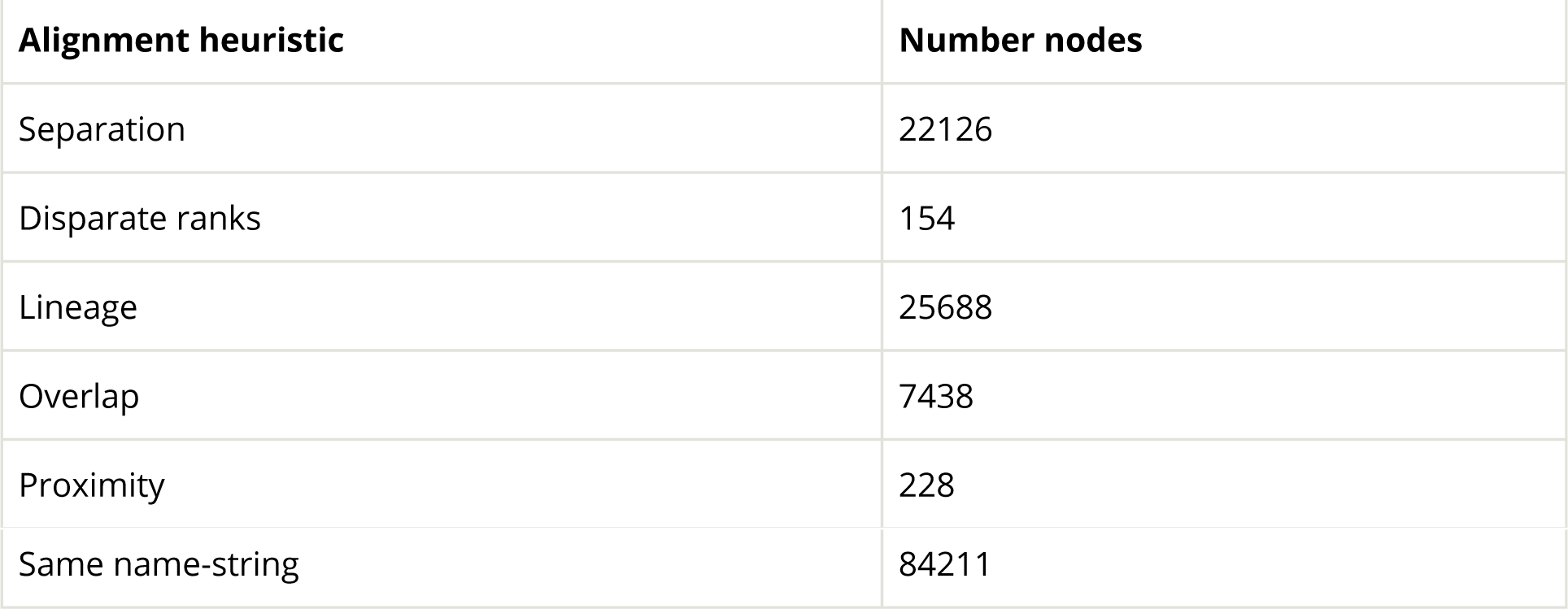
Download as CSV

Frequency of success of alignment heuristics. In cases where there were multiple candidate nodes, this table lists the number of times that a particular heuristic was the one to select a single candidate. Heuristics are listed in the order in which they are applied. Success of an ealier heuristics means that a later heuristic is not used for a given node.

After assembly, the next step in the method is to merge the unaligned nodes into the workspace taxonomy. Of the 3,780,949 unaligned nodes, the vast majority (99%) are grafted into the workspace. The remaining nodes (<1%) are either insertions, absorptions or remain unmerged due to ambiguities.

We also examined the fate of nodes from each of the input taxonomies, and Table 4 provides these results. The results are dependent on the order in which sources are added to the workspace. Overall, the number of conflicts is relatively low (<1 %).

**Table 4.**
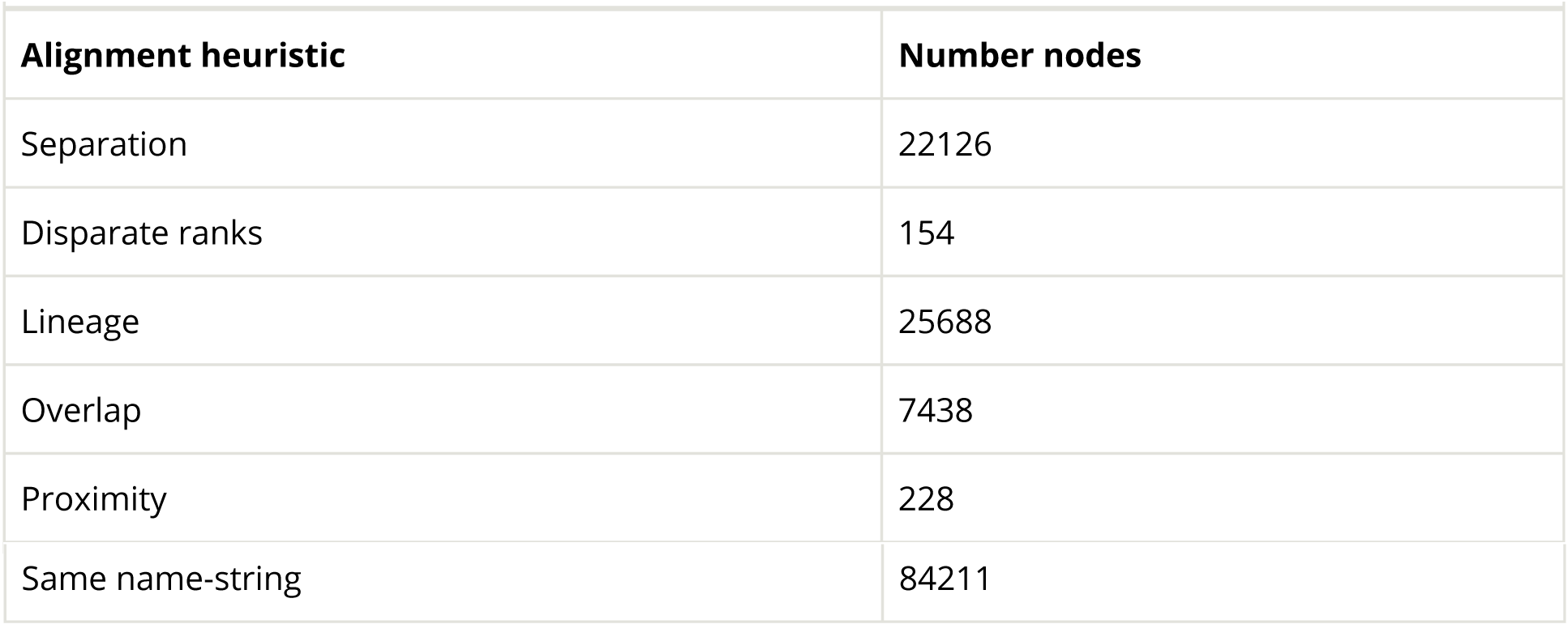
Download as CSV. Fate of source nodes from each of the input taxonomies. Unaligned nodes are either copied into the workspace or absorbed. Aligned nodes are added to the workspace through grafting or insertion.

## Evaluating the taxonomy relative to requirements

The introduction sets out requirements for an Open Tree taxonomy. How well are these requirements met?

### OTU coverage

We set out to cover the OTUs in the Open Tree corpus of phylogenetic trees. The corpus contains published studies (each study with one or more phylogenetic trees) that are manually uploaded and annotated by Open Tree curators. The user interface contains tools that help curators map the OTUs in a study to taxa in OTT. Of the 3,242 studies in the Open Tree database, 2,871 have at least 50% of OTUs mapped to OTT. (A lower overall mapping rate usually indicates incomplete curation, not an inability to map to OTT.) These 2,871 studies contain 538,728 OTUs, and curators have mapped 514,346 to OTT taxa, or 95.5%.

To assess the reason for the remaining 4.5% of OTUs being unmapped, we investigated a random sample of ten OTUs. In three cases, the label was a genus name in OTT followed by “sp” (e.g. *“Euglena* sp”), suggesting the curator’s unwillingness to take the genus as the correct mapping for the OTU. In the remaining seven cases, the taxon was already in OTT, and additional curator effort would have found it. Two of these were misspellings in the phylogeny source; one was present under a slightly different name-string (subspecies in OTT, species in study, the study reflecting a very recent reclassification); and in the remaining four cases, either the taxon was added to OTT after the study was curated, or the curation task was left incomplete. None in the sample reflected a coverage gap.

Of the 194,100 OTT records that are the targets of OTUs, 188,581 (97.2%) are represented in NCBI Taxonomy. If the Open Tree project had simply adopted NCBI Taxonomy instead of OTT, it would have met its OTU coverage requirement (but not the taxonomic coverage requirement). By comparison, GBIF covers 87.6%, and IRMNG covers 62.8%. The high coverage by NCBI reflects a preference among our curators for molecular phylogenetic evidence over other kinds.

### Phylogenetically informed classification

Assessing whether OTT is more ‘phylogenetically informed’ than it otherwise might be is difficult. The phylogenetic quality of the taxonomy is determined from by the taxonomic sources and their priority order. We have relied on the project’s curators, who have a strong phylogenetic interest, to provide guidance on both. Following are examples of curator decision-making:

- For microbes, SILVA is considered more phylogenetically sound than NCBI taxonomy, because the SILVA taxonomy is based on a recent comprehensive phylogenetic analysis.
- Priority of NCBI Taxonomy over the GBIF backbone is suggested by NCBI’s apparent interest in phylogeny, reflected in NCBI Taxonomy’s much higher resolution, its inclusion of phylogenetically important non-Linnaean groups such as Eukaryota, and by its avoidance of known paraphyletic groupings such as Protozoa.
- The Hibbett 2007upper fungal taxonomy reflects, by construction, results from the most recent phylogenetic studies of Fungi.

Ideally we would have a measure of ‘phylogenetically informed’ that we could use to compare OTT to other taxonomies, to test alternative constructions of OTT, and to check the forward progress of OTT. It is not clear what one would use as a standard against which to judge. The Open Tree project’s supertree of life is a candidate, but is not without issues (such as its own possible errors, and the fact that OTT is itself use in construction the supertree). Ensuring that comparisons are meaningful, and comparable with one another, would be a technical challenge.

### Taxonomic coverage

OTT has 2.3M binomials (presumptive valid species names), vs. 1.6M for Catalogue of Life (CoL). Since the GBIF source we used includes the Catalogue of Life, OTT includes all species in CoL.The number is larger in part because the combination of the inputs has greater coverage than CoL, and in part because OTT has many names that are either not valid or not currently accepted.

This level of coverage would seem to meet Open Tree’s taxonomic coverage requirement as well as any other available taxonomic source.

### Ongoing update

We aimed for a procedure that would allow simple re-building from sources, and also easy incorporation of new versions of sources. Re-building OTT version 3.0 from sources requires 17 minutes of real time. Our process currently runs on a machine with 16GB of memory; 8GB is not sufficient.

In the upgrade from 2.10 to 3.0, we added new versions of both NCBI and GBIF. NCBI updates frequently, so changes tend to be manageable and incorporating the new version was simple. In contrast, the version from GBIF represented both a major change in their taxonomy synthesis method. Many taxa disappeared, requiring changes to our ad hoc patches during the normalization stage. In addition, the new version of GBIF used a different taxonomy file format, which requires extensive changes to our import code (most notably, handling taxon name-strings that now included authority information).

We estimate the update from OTT 2.10 to OTT 3.0 required approximately three days of development time related to source taxonomy changes. This was greater than previous updates due to the changes required to handle the major changes in GBIF content and format.

### Open data

As the Open Tree project did not enter into any data use agreements in order to obtain OTT’s taxonomic sources, it is not obliged to require any such agreement from users of OTT. (A data use agreement is sometimes called ‘terms of use’. Legally, a DUA is a kind of contract). Therefore, users are not restricted in this way. In addition, the taxonomy is not creative expression, so copyright controls do not apply (Patterson et al. 2014). Therefore, to the best of our knowledge, use of OTT is unrestricted.

## Discussion

The primary actionable information in the source taxonomies consists of name-strings, and therefore the core of our method is a set of heuristics that can handle the common problems encountered when trying to merge hierarchies of name-strings. These problems include expected taxonomic issues such as synonyms, homonyms, and differences in placement and membership between sources. They also include errors such as duplications, spelling mistakes, and misplaced taxa. The problem cases add up to over 100,000 difficult alignments when the total number of source records measures over 6 million.

Ultimately there is no fully automated and foolproof test to determine whether two nodes can be aligned - whether node A and node B, from different source taxonomies, are about the same taxon. The information to do this is in the literature and in databases on the Internet, but often it is (understandably) missing from the source taxonomies.

It is not feasible to investigate such problems individually, so the taxonomy assembly methods identify and handle thousands of ‘special cases’ in an automated way. We currently use only name-strings, rudimentary classification information, and (minimally) ranks to guide assembly. We note the large role that our hand-curated “separation taxonomy” played in the alignment phase. This is a set of taxa that are consistent across the various sources, and allow us to make the (seemingly obvious) determination “these two taxa are in completely separate groups, so do not align them”.

## Open Tree Taxonomy as a taxonomy

We have developed the Open Tree Taxonomy (OTT) for the very specific purpose of aligning and synthesizing phylogenetic trees. We do not intend it to be a reference for nomenclature, or to substitute for expert-curated taxonomic databases. Several features of OTT make it unsuitable for taxonomic and nomenclatural purposes. It contains many names that are either not valid or not currently accepted. Some of these come from DNA sequencing via NCBI Taxonomy, which is also not a taxonomic reference, while others come directly from phylogenies submitted by Open Tree curators via our taxonomy curation features. OTT also contains more homonyms as compared to its sources. Most of these are duplicates that are artifacts of the assembly heuristics. For our purposes, these are not of great concern - when mapping OTUs in trees to taxa in OTT, we generally restrict mapping to a specific taxonomic context, and if there are multiple matches to OTT taxa with the same name, a curator can clearly see this situation and choose the taxon with the correct lineage.

## Community curation

We have also developed a system for curators to directly add new taxon records to the taxonomy from published phylogenies, which often contain newly described species that are not yet present in any source taxonomy. These taxon records include provenance information, including references explaining the taxon, and the identity of the curator. We expose this provenance information through the web site and the taxonomy API.

We also provide a feedback mechanism on the synthetic supertree browser, and find that most of the comments left are about taxonomy. Expanding this feature to capture this feedback in a more structured, and therefore machine-readable, format would allow users to directly contribute taxonomic patches to the system.

## Comparison to other taxonomies

Given the very different goals of the Open Tree Taxonomy in comparison to most other taxonomy projects, it is difficult to compare OTT to other taxonomies in a meaningful way. The Open Tree Taxonomy is most similar to the GBIF taxonomy, in the sense that both are a synthesis of existing taxonomies rather than a curated taxonomic database. The GBIF method is yet unpublished (for basic information on the GBIF backbone see Döring 2016a,Döring 2016b). Once the GBIF method has been formally described, it will be useful to compare the two approaches and identify common and unique techniques for automated, scalable name-string matching.

## Potential improvements and future work

The development of the assembly software has been driven by the needs of the Open Tree project, not by any concerted effort to create a widely applicable or theoretically principled tool. Many improvements are possible on both practical and theoretical grounds. Following are some of the directions for development that could have the highest impact.

- It is very likely that alignment could be improved by making better use of species proximity implied by the shape of the classification, and decreasing its reliance on the names of internal nodes. Better use of proximity might permit separation and identification of tips without use of a separation taxonomy, removing the need for the manual work of maintaining the separation taxonomy and the adjustment directives needed to align source taxonomies to it.
- Additional information that is available in some source taxon records could be put to good use in alignment, especially authority information. Names could also be analyzed to detect partial matches, e.g. matching on species epithets even when the genus disagrees, and spelling and gender variant recognition. Other work on name matching (Patterson et al. 2016) goes far beyond what is done for OTT and these techniques should be used.
- An assembly run can lead to a variety of error conditions and test failures. Currently these are difficult to diagnose, mainly for lack of technology for displaying the particular pieces of the sources, workspace, and assembly history that are relevant to the error. Once this information is surfaced it is usually not too difficult to work out a fix in the form of a patch or an improvement to the program logic.
- The community curation should be developed, as mentioned above. Its success would depend on allowing users to test proposed changes and diagnose and repair any problems with them.
- Curators frequently request new taxonomy sources. The most frequently requested are improved fish, bird, and plant sources. Again, the information is generally available, but not yet harvested. (Some frequently requested sources may only be accessed under agreement with contractual terms (variously called “terms of use” or a “data use agreement”). One of these is the IUCN Red List (International Union for Conservation of Nature and Natural Resources 2016), an important source of up-to-date information on mammal species. These sources are off limits to Open Tree due to the project’s open data requirement.)
- The presence of invalid and unaccepted names remains a significant problem. The information needed to detect them is available, and could be harvested.
- Basic usability features for application to new projects would include proper packaging of the application, and support for Darwin Core (Wieczorek et al. 2012).

Future work on taxonomy aggregation should attempt a more rigorous and pluralistic approach to classification (Franz et al. 2016). Alignment should detect and record lumping and splitting events, and the classification conflicts detected during merge should be exposed to users. Exposing conflicts is in the interest of scientific transparency. Retaining alternative groupings could be useful in phylogenetic analysis, as a check on which of the sources agree or disagree with a given analysis. Lumping and splitting, when they can be detected, could be recorded as taxon that has, as one of its children, a distinct taxon with the same name-string. Ideally better handling of “taxon concepts” in aggregators would encourage sources to make links to primary sources more readily available for a variety of purposes.

## Data resources

All source code is open source (licensed BSD 2-clause) and available on GitHub at https://github.com/OpenTreeOfLife/reference-taxonomy. A snapshot of the code used to produce the version of OTT described here is archived at Zenodo (link needed). All data, including Open Tree Taxonomy 3.0 and all processed source taxonomies is archived on Dryad (cannot be uploaded before paper accepted; version for review at https://github.com/OpenTreeOfLife/reference-taxonomy/tree/master/doc/method/data-package).

## Conclusions

We have presented a method for merging multiple taxonomies into a single synthetic taxonomy. The method is designed to produce a taxonomy optimized for the Open Tree of Life phylogeny synthesis project. Most taxonomy projects are databases of taxonomy information that are continuously updated by curators as new information is published in the taxonomic literature. In contrast, the Open Tree Taxonomy takes several of these curated taxonomies and assembles a synthetic taxonomy *de novo* each time a new version of the taxonomy is needed.

We have also developed a system for curators to directly add new taxa to the taxonomy from published phylogenies. These taxon additions include provenance information, including the source of the taxon and identity of the curator. We expose this provenance information through the website and the taxonomy API. Most of the Open Tree feedback has been about taxonomy, and expanding this feature to other types of taxonomic information allows users to directly contribute expertise and allows projects to easily share that information.

Taxonomic information is certainly best curated at a scale smaller than “all life” by experts in a particular group. Therefore, producing comprehensive taxonomies should be a synthesis of curated taxonomies. We advocate for the type of methods being used by Open Tree and by GBIF, where synthesis is done in a repeatable fashion from sources, allowing changed information in sources to be quickly included in the comprehensive taxonomy. Provenance information is retained and presented as part of the synthetic taxonomy. This type of synthesis requires that source taxonomies be available online, either through APIs or by bulk download, in a format that can be easily parsed, and ideally without terms of use that prevent distribution and reuse of the resulting synthetic taxonomies.

## Acknowledgements

We would like to thank Nico Franz, Mark Holder, and Alan Ruttenberg for their comments on early versions of this manuscript;Markus Döring, Tony Rees, and Paul Kirk for answering our many questions about source taxonomies; Cody Hinchliff for writing an early version of the assembly code and etablishing the general approach to taxonomy combination; and Yan Wong and other users of the Open Tree system provided many helpful comments on the taxonomy.

## Funding program

NSF AVAToL (Assembling Visualizing, and Analyzing the Tree of Life) DEB-1208809

## Grant title

Automated and community-driven synthesis of the tree of life

## Hosting institution

Duke University

## Author contributions

JAR designed and implemented the taxonomy assembly system; KAC and JAR wrote the paper.

